# Imputed DNA methylation outperforms measured loci associations with smoking and chronological age

**DOI:** 10.1101/2024.09.05.611501

**Authors:** Anne Richmond, Jure Mur, Sarah E Harris, Janie Corley, Hannah R Elliott, Christopher N Foley, Eilis Hannon, Zhana Kuncheva, Josine L Min, Mahdi Moqri, Magatte Ndiaye, Benjamin B Sun, Catalina A Vallejos, Kejun Ying, Vadim N Gladyshev, Simon R Cox, Daniel L McCartney, Riccardo E Marioni

## Abstract

Multi-locus signatures of blood-based DNA methylation are well-established biomarkers for lifestyle and health outcomes. Here, we focus on two CpGs that are strongly associated with age and smoking behaviour. Imputing these loci via epigenome-wide CpGs results in stronger associations with outcomes in external datasets compared to directly measured CpGs. If extended epigenome-wide, CpG imputation could augment historic arrays and recently-released, inexpensive but lower-content arrays, thereby yielding better-powered association studies.

## Main

Over the past decade, Illumina array technology has led the profiling of DNA methylation (DNAm) in large cohort studies^1,2^. This began with the measurement of ~27,000 CpG loci (27K array), followed by ~450,000 (450K array) and most recently 800,000+ CpG sites (EPICv1 and EPICv2). The vast majority of content on the smaller 27K and 450K arrays are also contained on the EPIC arrays^3,4^. By leveraging the widespread correlations across the methylome^5,6^, it may be possible to derive an imputation framework to augment existing datasets with missingness or those that were generated using historic arrays.

The requirement for such a tool is further emphasised by the recent development and launch of the Illumina Methylation Screening Array (MSA)^7^. The MSA is the most affordable DNAm array to date and was designed specifically for application in large biobank studies. The MSA array assesses DNAm at 269,094 sites of which 145,318 are present on the EPICv2 array^8^. The MSA therefore contains new content with enrichment for sites linked to regulatory and cell-specific chromatin states^8^. Clearly, this will create major challenges when trying to replicate findings or meta-analyse data across array technologies. However, if one can identify ways to impute content, this will not only benefit cohorts with existing data, but will also afford an opportunity to assess DNAm at greater scale, via a less expensive method, prior to boosting content through imputation. Finally, given that imputation considers CpG information from multiple loci, this averaging process may lead to fewer spurious outliers at well-imputed sites, resulting in better-powered association studies and improved multi-CpG biomarkers, such as epigenetic clocks^9^.

Building an accurate DNAm imputation server is therefore of immense value to the research community. However, developing this tool is non-trivial both in terms of scale (~1 million unique CpG sites to consider) and determining imputation quality for traits that are both dynamic and influenced by multiple factors.

To highlight the potential of the work, we present pilot findings for two CpG loci that are established blood-based correlates of chronological age (cg16867657, *ELOVL2*)^10,11^ and smoking behaviour (cg05575921, *AHRR*)^12,13^.

Similar to polygenic risk scores, methylation-based predictors can be derived as linear, weighted additive combinations of CpGs. Hereafter, we refer to these as Epigenetic Scores (EpiScores). EpiScores for the two CpGs were derived using data from 18,869 volunteers from the Generation Scotland cohort. DNAm was profiled from blood samples collected between 2006 and 2011, when individuals were aged between 17 and 99 years (11,098, 58.8% female, **Supplementary Figure 1**). The EPICv1 array was used to measure DNAm – full details of the processing and quality control are presented elsewhere^14^ and briefly summarised in **Online Methods**. There were methylation estimates available for 752,722 CpGs after quality control. These were subset to loci present on the 450k array to maximise backwards compatibility, and further filtered to the target locus (cg16867657 or cg05575921) and the 200,000 most variable probes (after excluding the target CpG) for computational efficiency and to remove invariant CpG sites. In a final quality control step, each CpG M-value was pre-adjusted for sex, analysis batch and the first 10 genetic principal components^15^ via linear regression in R with the resulting residuals taken forward for the main analysis.

Elastic net penalised linear regression was used to derive EpiScores using the *biglasso* package (version 1.3.7)^16^ in R (version 4.0.3). The target CpG (cg16867657) was specified as the outcome variable with the 200,000 most variable CpGs as the predictors. This was then repeated with cg05575921 as the outcome. 20-fold cross-validation was applied to obtain the optimal lambda (shrinkage parameter) that minimised the mean error. The subsequent models resulted in 65 non-zero coefficients for both cg16867657 and cg05575921. These coefficients are presented in **Supplementary Table 1**.

The predictors were then tested in external datasets. The age-related CpG EpiScore was tested in the publicly available dataset used by Hannum et al.^17^ to derive one of the first epigenetic clocks (GSE40279, 450K array). After downloading the data (n=656 individuals aged 19 to 101 years), the model weights were applied (65/65 CpGs present) and the EpiScore was derived. The measured CpG and CpG EpiScore were highly correlated with each other (Pearson r = 0.90, P = 8.1×10^−238^). **Figure 1** shows that the imputed CpG EpiScore yielded a stronger, more significant correlation with chronological age than the measured CpG: Pearson r_EpiScore_ = 0.88 (P = 8.4×10^−214^) versus r_cg16867657_ = 0.83 (P = 7.4×10^−167^). A second publicly available dataset (GSE246337^18^, EPICv2, 59/65 CpGs available), which contained 500 individuals from the Mass General Brigham (MGB) Biobank was also considered. This cohort was evenly divided by sex and representing ages 18-99 years and a broad range of ethnicities. The dataset was subset to 437 individuals with no missing data for age or the 59 CpGs. Here, the CpG – EpiScore correlation was 0.96 (P = 2.0×10^−239^) with age correlations of r_EpiScore_ = 0.94 (P = 3.7×10^−204^) and r_cg16867657_ = 0.93 (P = 1.6×10^−185^). When the cohort was stratified into age decades, the EpiScore – CpG correlation decreased (r_range_ = 0.64-0.81) but remained larger than the within-strata associations between either variable and age (**Supplementary Table 2**).

**Figure 1.**
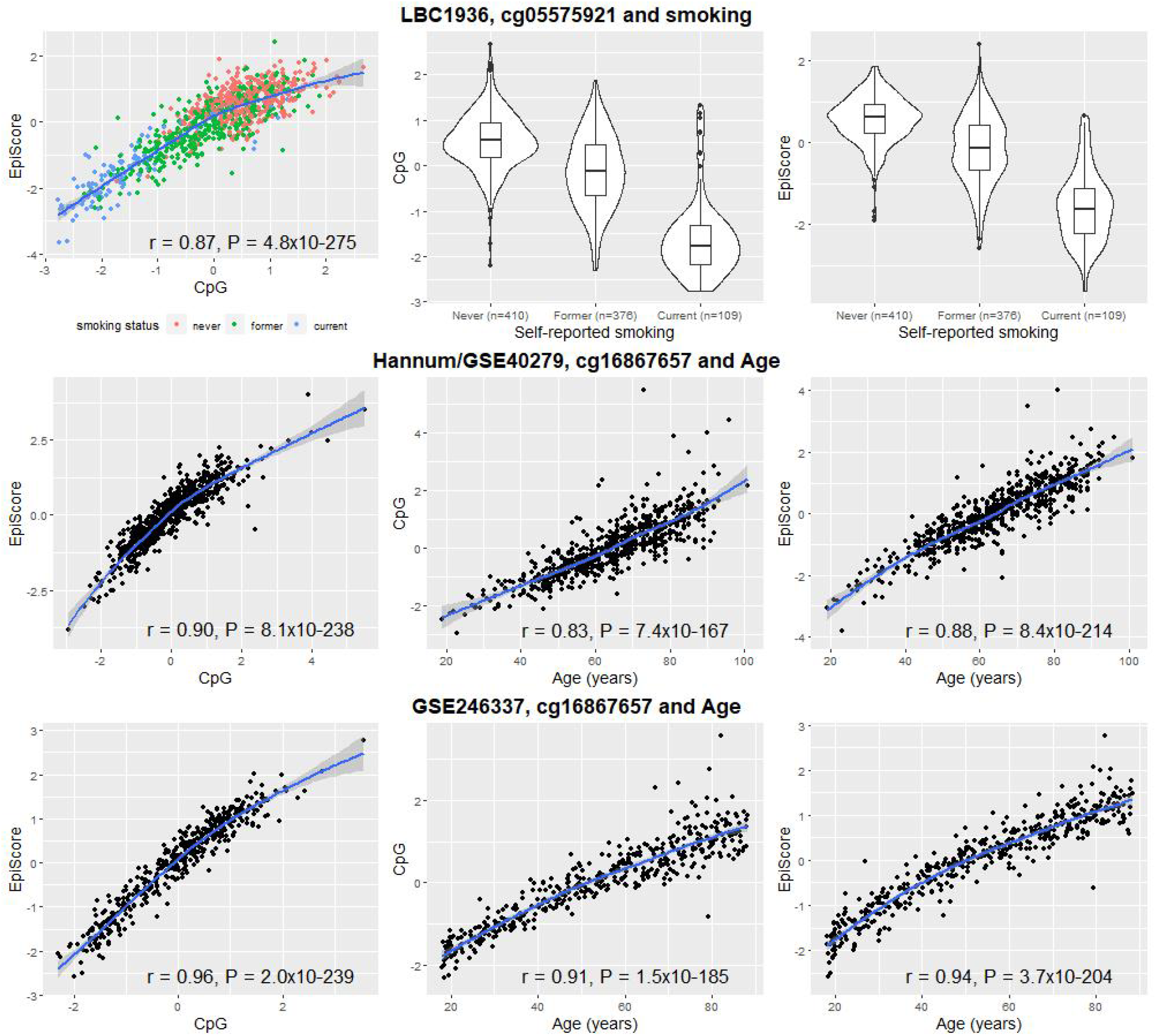
Comparison of measured versus imputed CpGs for smoking (top row) and age (bottom rows) and their associations with smoking (Lothian Birth Cohort 1936, n = 895) and age (‘Hannum’ dataset, GSE40279, n= 656; GSE246337, n=500). Measured/imputed CpGs were scaled to mean 0, variance 1. Blue lines denote local regression lines (loess estimation) with 95% confidence intervals.

The smoking-related CpG EpiScore was tested in 895 individuals from the Lothian Birth Cohort 1936^19^. All individuals were born in 1936 and had a mean age of 70 years (SD = 0.8) with 442 (49.4%) females when blood samples were obtained. DNAm was profiled from these samples using the Illumina 450K array^20^. Details of quality control are presented in the **Online Methods**. Smoking pack years information was obtained by self-report questionnaires and calculated as years smoked (age stopped minus age started smoking) multiplied by the number of 20-cigarette packs smoked per day. This information was available for 881 of the 895 participants and underwent a log(pack years + 1) transformation to reduce skew. 64/65 CpGs were available in LBC1936 for the projection of the EpiScore.

The Pearson correlation between the measured cg05575921 and its EpiScore was r = 0.87 (P = 4.8×10^−275^) in the whole population and r = 0.61 in the sub-group of n = 410 never-smokers. This is plotted in **Figure 1** alongside boxplots of the measured CpG and EpiScore against self-reported smoking status. The measured CpG and its EpiScore associations with smoking pack years were r = −0.62 (P = 8.8×10^−94^) and r = −0.60 (P = 2.4×10^−87^), respectively. The EpiScore also showed a better classification of current versus never smokers (assessed by self-report, n = 109 and 410, respectively) in the same population: area under the receiver operating characteristic curve of 0.971 versus 0.954. A sensitivity analysis training the EpiScores using DNAm beta-values in place of M-values made minimal differences to the results (**Supplementary Table 3**).

Together, these findings show that imputation of CpG methylation from other CpG sites leads to stronger and more statistically significant associations with two important outcomes for health research: age and smoking. While the imputation success at the selected sites is part-driven by their well-established associations with age and smoking, these findings militate for further work to assess how well the approach generalises across all CpGs present on Illumina arrays. In addition, family-structure/relatedness was not accounted for within the Generation Scotland training cohort, which may have led to information leakage across folds and overfitting. However, we tested the resulting EpiScores in external datasets where the DNAm was also processed and normalised independently. Further tests need to be carried out to ensure that the resulting signatures translate across diverse populations. Here, the LBC test cohort contained individuals of Scottish ancestry while the Hannum dataset contained a mixture of European and Hispanic ancestry individuals (n = 426 and 230, respectively) and GSE246337 contained a mix of European-, African-, Asian-, and Hispanic-ancestry individuals. Subsetting to CpGs that are commonly found on the 450K, EPIC and MSA arrays prior to training EpiScores would maximise the gains for all cohorts. Further subsetting this list to loci that have similar patterns (e.g., mean and SD by age and sex) across populations, as well as exploring the properties of well-imputed sites (e.g., by genomic location or SNP-based heritability) will further inform the generalisability of the findings. Future studies should also focus on incorporating genotypic contributions to CpG variability^21^ or more flexible imputation approaches that can capture non-linear patterns.

To further highlight the utility of our approach, we applied a similar methodology to predict GDF15 protein levels from other plasma proteins in UK Biobank (**Online Methods**). The imputed variable correlated 0.87 with measured GDF15 in the test set and showed a larger and more statistically significant association (per SD) with incident dementia (n_cases_ = 590, n_controls_ = 17,952): Hazard Ratio (HR)_GDF15_ = 1.37, 95% CI [1.26, 1.48], P = 3.1×10^−15^ and HR_GDF15_ProteinScore_ = 1.42 [1.31, 1.54], P = 2.5×10^−17^.

In conclusion, the imputation of array-based CpG methylation and plasma proteins is feasible and can lead to larger and more statistically significant effect sizes in association studies for complex traits.

## Availability of data and material

According to the terms of consent for Generation Scotland participants, access to data must be reviewed by the Generation Scotland Access Committee. Applications should be made to.

Further information on accessing Lothian Birth Cohort data from the Lothian Birth Cohort Study, University of Edinburgh is available online, alongside a data request form and data dictionaries (https://www.ed.ac.uk/lothian-birth-cohorts/data-access-collaboration).

Data from UK Biobank is available to approved researchers directly from UK Biobank (https://www.ukbiobank.ac.uk/enable-your-research/apply-for-access).

## Acknowledgements

This research was funded in whole, or in part, by the Wellcome Trust (104036/Z/14/Z and 221890/Z/20/Z). For the purpose of open access, the author has applied a CC BY public copyright license to any Author Accepted Manuscript version arising from this submission. Generation Scotland received core support from the Chief Scientist Office of the Scottish Government Health Directorates (CZD/16/6) and the Scottish Funding Council (HR03006). DNA methylation profiling of the Generation Scotland samples was carried out by the Genetics Core Laboratory at the Edinburgh Clinical Research Facility, Edinburgh, Scotland, and was funded by the Medical Research Council UK and Wellcome (Wellcome Trust Strategic Award STratifying Resilience and Depression Longitudinally (STRADL; Reference 104036/Z/14/Z). DNA methylation data for Generation Scotland was also funded by a 2018 NARSAD Young Investigator Grant from the Brain & Behavior Research Foundation (Ref: 27404; awardee: Dr David M Howard) and by a John, Margaret, Alfred and Stewart Sim Fellowship from the Royal College of Physicians of Edinburgh (Awardee: Dr Heather C Whalley).

We thank the LBC1936 participants and team members who contributed to these studies. The LBC1936 is supported by the Biotechnology and Biological Sciences Research Council, and the Economic and Social Research Council [BB/W008793/1] (which supports S.E.H., and J.C.), Age UK (Disconnected Mind project), the Milton Damerel Trust, the Medical Research Council (G0701120, G1001245, MR/M013111/1, MR/R024065/1) and the University of Edinburgh. Methylation typing of LBC1936 was supported by the Centre for Cognitive Ageing and Cognitive Epidemiology (Pilot Fund award), Age UK, The Wellcome Trust Institutional Strategic Support Fund, The University of Edinburgh, and The University of Queensland.

The research was conducted using the UK Biobank resource, with approved project number 10279.

H.R.E. and J.L.M. are members of the Medical Research Council Integrative Epidemiology Unit at the University of Bristol, which is supported by the Medical Research Council and the University of Bristol (MC_UU_00032/3, MC_UU_00032/4 and MC_UU_00032/5). K.Y. is supported by NIA F99/K00 grant FAG088431A. This research was also supported in part by NIA and Hevolution grants (awardee: Vadim Gladyshev). S.R.C. was supported by a Sir Henry Dale Fellowship jointly funded by the Wellcome Trust and the Royal Society (Grant Number 221890/Z/20/Z). J.M. and R.E.M. are supported by Alzheimer’s Society project grant AS-PG-19b-010.

## Ethics approval and consent to participate

All components of GS received ethical approval from the NHS Tayside Committee on Medical Research Ethics (REC Reference Number: 05/S1401/89). GS has also been granted Research Tissue Bank status by the East of Scotland Research Ethics Service (REC Reference Number: 20-ES-0021), providing generic ethical approval for a wide range of uses within medical research.

Ethical approval for the LBC1936 study was obtained from the Multi-Centre Research Ethics Committee for Scotland (Wave 1, MREC/01/0/56) and the Lothian Research Ethics committee (Wave 1, LREC/2003/2/29).

All participants provided written informed consent. These studies were performed in accordance with the Helsinki declaration.

## Online Methods

Tabular overview of the DNA methylation collection and QC process for Generation Scotland and the Lothian Birth Cohort 1936.

|  | Generation Scotland | Lothian Birth Cohort 1936 |
| --- | --- | --- |
| **Tissue** | Blood | Blood |
| **N** | 18,869 | 895 |
| **Array** | EPICv1 | 450k |
| **Comments** | Processed in 4 sets (n=5,087, 4,450, 8,876, and 456) | Processed in 2 sets (n=861 and 34) |
| **Background correction** | Background and dye-bias correction were carried out using meffil<sup>1</sup> (versions 1.1.0 and 1.1.2) and the "noob" method | Raw intensity data were background-corrected and normalised using internal controls |
| **Probe removal** | (i) outliers from visual inspection of the log median intensity of the methylated versus unmethylated signal per array,<br>(ii) beadcount < 3 in more than 5% of samples,<br>(iii) $\geq 5\%$ of samples having a detection p-value > 0.05,<br>(iv) annotated to X or Y chromosome,<br>(v) overlapped with SNPs<br>(vi) potentially cross-hybridizing | (i) low (< 95%) detection rate with p-value < 0.01,<br>(ii) inadequate hybridization, bisulfite conversion, nucleotide extension, or staining signal (manual inspection) |
| **Sample removal** | (i) sex mismatch,<br>(ii) $\geq 1\%$ of CpGs with detection p-value > 0.05,<br>(iii) if sample was not blood-based | (i) low call rate (<450,000 CpGs at P<0.01),<br>(ii) sex mismatch<br>(iii) samples exhibiting a poor match between genotype and SNP control probes |
| **Normalisation** | Dasen<sup>2</sup> | Danet<sup>2</sup> |
| **N CpGs post QC** | 752,722 | 450,276 |

## GDF15 Analysis

### UK Biobank Overview

UK Biobank is a prospective cohort study of around 500,000 individuals living across England, Wales and Scotland^3^. Participants were aged between 40 and 70 years at recruitment between 2006 and 2010. Demographic, lifestyle, and physiological data were collected via self-report and at a baseline clinic visit. Blood samples were obtained from the majority of volunteers, most of whom provided consent for linkage to their electronic primary and secondary care health records.

### Protein Measurement

Proteomic data were generated for 53,014 participants using the Olink Explore panel, which assesses 2,923 proteins (data field 30900)^4^. Proteins with >20% missingness (n=12), participants missing >20% of the 2,923 proteins (n = 8,834), and participants missing measurements for GDF15 (n = 865) were excluded.

### Dementia Ascertainment

Dementia was ascertained using the algorithmically-defined outcomes in UK Biobank (data field 42018). These combine data from self-report at baseline, hospital records, and death register records to increase the positive predictive value of dementia ascertainment^5^.

### Statistical Analyses

The cohort with available protein data was split at random into equally-sized training and test sets (n_train_ = 21,888, n_test_ = 21,889). Each set was then processed separately. First, missing values for all proteins apart from GDF15 were mean imputed. Second, proteins were scaled to have mean zero and unit variance.

Elastic net penalised linear regression was used to derive a ProteinScore for GDF15 using the *biglasso* package (version 1.3.7)^6^ in R (version 4.0.3). GDF15 was specified as the outcome variable with the 2,911 remaining proteins listed as the predictors. An elastic net penalty (alpha = 0.5) was set and 20-fold cross-validation was applied to obtain the optimal lambda (shrinkage parameter) that minimised the mean error. The subsequent model resulted in 1,281 non-zero coefficients plus an intercept. These coefficients are presented in **Supplementary Table 4** and were used to generate a ProteinScore for GDF15 in the test set.

A Pearson correlation was used to compare measured GDF15 with its ProteinScore in the test set. This was followed by Cox proportional hazards regression analyses, adjusting for age and sex (data fields 34, 52, 53 and 31) and either GDF15 or its ProteinScore as the predictor of interest. Right-censoring occurred when a person was diagnosed with dementia, died (data field 40000), was lost to follow-up (data field 191), or on the last day of data availability (31^st^ December 2022), whichever came first. To exclude individuals too young to experience late-onset dementia, the sample was restricted to participants born before the year 1962. In the test set, this resulted in 590 individuals with an incident dementia diagnosis and 17,952 controls.

---

**Supplementary Figure 1.**
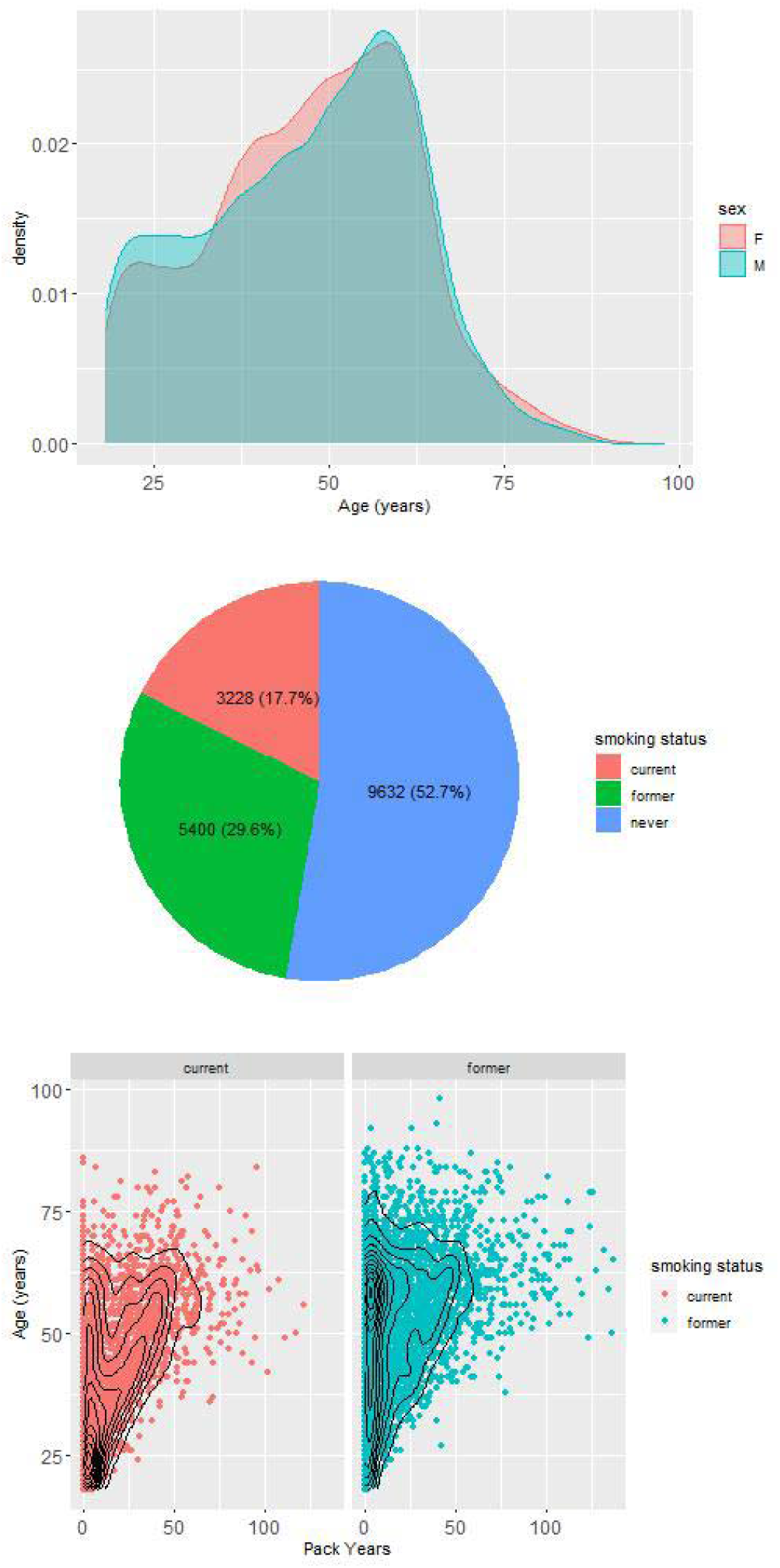
Overview of the basic study demographics for Generation Scotland. The top panel shows the distribution of age by sex with males in turquoise and females in light red. The middle panel shows the distribution of smoking status and the lower panel shows a scatter plot with density contours for number of pack years of smoking by age for both current (light red) and former (turquoise) smokers.

